# Spatial relations trigger visual binding of people

**DOI:** 10.1101/2020.10.01.322198

**Authors:** Parvaneh Adibpour, Jean-Rémy Hochmann, Liuba Papeo

**Author notes:** **Corresponding author:** Parvaneh Adibpour, CNRS, Institut des Sciences Cognitives - Marc Jeannerod, 67 Boulevard Pinel, 69675, Bron, France.

## Abstract

To navigate the social world, humans must represent social entities, and the relationships between those entities, starting with spatial relationships. Recent research suggests that two bodies are processed with particularly high efficiency in visual perception, when they are in a spatial positioning that cues interaction, i.e. close and face-to-face. Socially relevant spatial relations such as facingness may facilitate visual perception by triggering grouping of bodies into a new integrated percept, which would make the stimuli more visible and easier to process. We used electroencephalography and a frequency-tagging paradigm to measure a neural correlate of grouping (or visual binding), while female and male participants saw images of two bodies face-to-face or back-to-back. The two bodies in a dyad flickered at frequency F1 and F2, respectively, and appeared together at a third frequency Fd (dyad frequency). This stimulation should elicit a periodic neural response for each body at F1 and F2, and a third response at Fd, which would be larger for face-to-face (*vs.* back-to-back) bodies, if those stimuli yield additional integrative processing. Results showed that responses at F1 and F2 were higher for upright than for inverted bodies, demonstrating that our paradigm could capture neural activity associated with viewing bodies. Crucially, the response to dyads at Fd was larger for face-to-face (*vs.* back-to-back) dyads, suggesting integration mediated by grouping. We propose that spatial relations that recur in social interaction (i.e., facingness) promote binding of multiple bodies into a new representation. This mechanism can explain how the visual system contributes to integrating and transforming the representation of disconnected body-shapes into structured representations of social events.

## Introduction

The visual world is an array of objects, but our understanding of the world goes beyond single object representation, involving, among many others, the processing of how individual entities relate to one another. The analysis of relations is essential to represent a scene or an event.

In social scenes (i.e., scenes involving multiple people), certain relational features, such as spatial proximity and positioning angle between bodies, are reliably correlated with the occurrence of social interaction (Zhou et al., 2019). These features are immediately available to our visual system, affecting the way in which individuals represent objects (Glanemann et al., 2016; Hafri et al., 2018; Papeo et al., 2017, 2019). For instance, it has been shown that, presented at perceptual threshold, two bodies are recognized more easily when they are shown face-to-face than when they are back-to-back (Bellot et al., 2021; Papeo et al., 2017). Face-to-face bodies are also attended to with higher priority, relative to other multiple-body configurations: in visual search through a crowd, face-to-face bodies recruit attention more strongly, and are searched for more efficiently, than back-to-back bodies (Papeo et al., 2019; Vestner et al., 2019, 2020)

Efficient processing of physically disconnected object-shapes in certain spatial configurations could be achieved through perceptual grouping, the binding of parts (here, bodies) into a new unitary representation (Coren & Girgus, 1980). Grouping based on spatial relations can reflect the gestalt laws of low-level vision (Koffka, 1935; Wagemans et al., 2012), but can also be driven by statistically frequent or semantically relevant relations (Kaiser et al., 2014; Kaiser et al., 2019) or, as suggested for face-to-face bodies, socially relevant spatial relations (Papeo, 2020; Quadflieg et al., 2015). In sum, grouping would give rise to a new composite representation, when the spatial arrangement of objects matches an expected, regular, familiar, or meaningful configuration. The processing of multiple objects as a coherent, unitary structure increases efficiency, possibly by reducing the stimulus complexity and/or the competition for selection between objects (Kaiser et al., 2019; McMains & Kastner, 2010; Reddy et al., 2009)

At the neural level, recent functional magnetic resonance imaging (fMRI) studies have reported that visual areas underlying body perception in the occipito-temporal cortex, participate in the representation of multiple-body stimuli, showing increased response and more accurate representation of dyads of face-to-face bodies (*vs*. non facing bodies or single bodies) (Abassi & Papeo, 2020; Bellot et al., 2020; Walbrin & Koldewyn, 2019). Activity in these areas is further modulated by the coherence of a multiple-person scene, differentiating between scenes depicting people that belong to the same (*vs.* a different) context (e.g., a club or a party; Quadflieg et al., 2015). Increased activity for socially relevant multiple-body configurations has been interpreted as evidence of grouping, following prior research on multiple-object perception. Indeed, research on multiple-object perception has consistently demonstrated an increase in the neural response to pairs of objects seen in familiar (i.e., regular or expected) configurations (e.g., a screen above a keyboard) as compared with the same objects in a spatial relation that does not promote the formation of composite representation (e.g., a screen below a keyboard; Kaiser et al., 2014; Kim & Biederman, 2011; MacEvoy & Epstein, 2009; Roberts & Humphreys, 2010). Increased activity for regular multiple-object configurations has been specifically associated with markers of integrative processing that could mediate grouping of multiple parts or multiple objects in a composite representation that is more than the sum of the parts (Baeck et al., 2013; Baldassano et al., 2016; Kaiser & Peelen, 2018; Kubilius et al., 2015).

Here, we sought to provide an objective measure of grouping of facing body dyads, using a frequency-tagging electroencephalography (EEG) paradigm to separate the response to the parts of a dyad (single bodies) from the response to the whole (the dyad). In frequency-tagging EEG, stimuli are presented periodically, at a regular frequency. The periodic stimulation entrains a periodic neural activity at the stimulation frequency, which is easily distinguishable in the frequency domain (Norcia et al., 2015; Regan, 1966). By presenting different parts of a multi-part stimulus at different stimulation frequencies, it is possible to dissociate the response to each single part from the response to the whole stimulus.

This paradigm has proven effective in capturing neural effects of binding visual features based on Gestalt principles (Aissani et al., 2011; Alp et al., 2016), and binding parts into an object representation (e.g., a face; Boremanse et al., 2013). In particular, using frequency-tagging EEG, Boremanse and colleagues (2013) could distinguish the responses to the two halves of a face, each presented at a different regular frequency, from the response to the whole face, emerging when the brain integrates information from the two halves, at a third regular frequency. Critically, the response to whole-faces was larger when the halves gave rise to a face, relative to when their arrangement violated the canonical face configuration (i.e., the halves were next to each other but misaligned; Boremanse et al., 2013). Thus, the response to whole-faces was not elicited by the mere occurrence of more facial features, but by the integration of those features in a well-formed face.

Extending the rationale of Boremanse et al. (2013), in the present study, we used a frequency-tagging paradigm to obtain objective responses to the parts and whole of a visual scene, where the constituting parts were two single bodies and the whole was the dyad. Each of the two bodies flickered at a different frequency to elicit a frequency-tagged response at the corresponding frequency (response to parts: F1 and F2). The two bodies appeared together in full view at a third frequency, that we called Fd (i.e., dyad-related frequency). To isolate visual activity associated with body perception, in half of the trials, bodies were presented inverted upside-down, a condition that significantly reduces activity in visual areas for body perception, while providing matched visual stimulation (Brandman & Yovel, 2010; Brandman & Yovel, 2016). In different trials, the two bodies faced toward or away from the center. Therefore, when they appeared together at Fd, they either faced toward each other (facing dyad) or away from each other (non-facing dyad). We reasoned that, if the face-to-face positioning gives rise to a new integrated representation of the two bodies, the response at Fd should be enhanced for facing (vs. non-facing) dyads. Enhanced response to facing dyads will provide initial evidence for multiple-body integration, triggered by specific –socially relevant– spatial relations. Grouping through integration may begin to account for the neural mechanism that transforms a group of bodies in the representation of a social event such as an interaction.

## Materials and Methods

### Participants

Twenty healthy adults (13 female; mean age 23.3, SD = 4.6) with normal or corrected-to-normal vision participated in the study. The sample size was decided based on following EEG studies with a similar paradigm (Alp et al., 2016; Boremanse et al., 2013; Kaiser et al., 2020). The study was approved by the local ethics committee. All participants gave written informed consent and received monetary compensation for participation.

### Stimuli

Gray-scale images of an identical body in six different postures, seen in profile view, were created with Daz3D (Daz Productions, Salt Lake City) and the Image Processing Toolbox of MATLAB (MathWorks). Four facing dyads (Figure 1a) were created by placing two bodies face-to-face. Four non-facing dyads were created by swapping the position of the two bodies in the facing dyads (Figure 1b). The distance between the center of each body and the center of the screen was fixed (77 pixels); furthermore, the distance between two bodies (i.e., between their inner edges on the horizontal axis) was matched between a facing dyad and its corresponding non-facing dyad (mean = 38.2 ± 5.4 pixels). Thus, the two bodies were at comparable distance from central fixation across facing and non-facing dyads.

**Figure 1.**
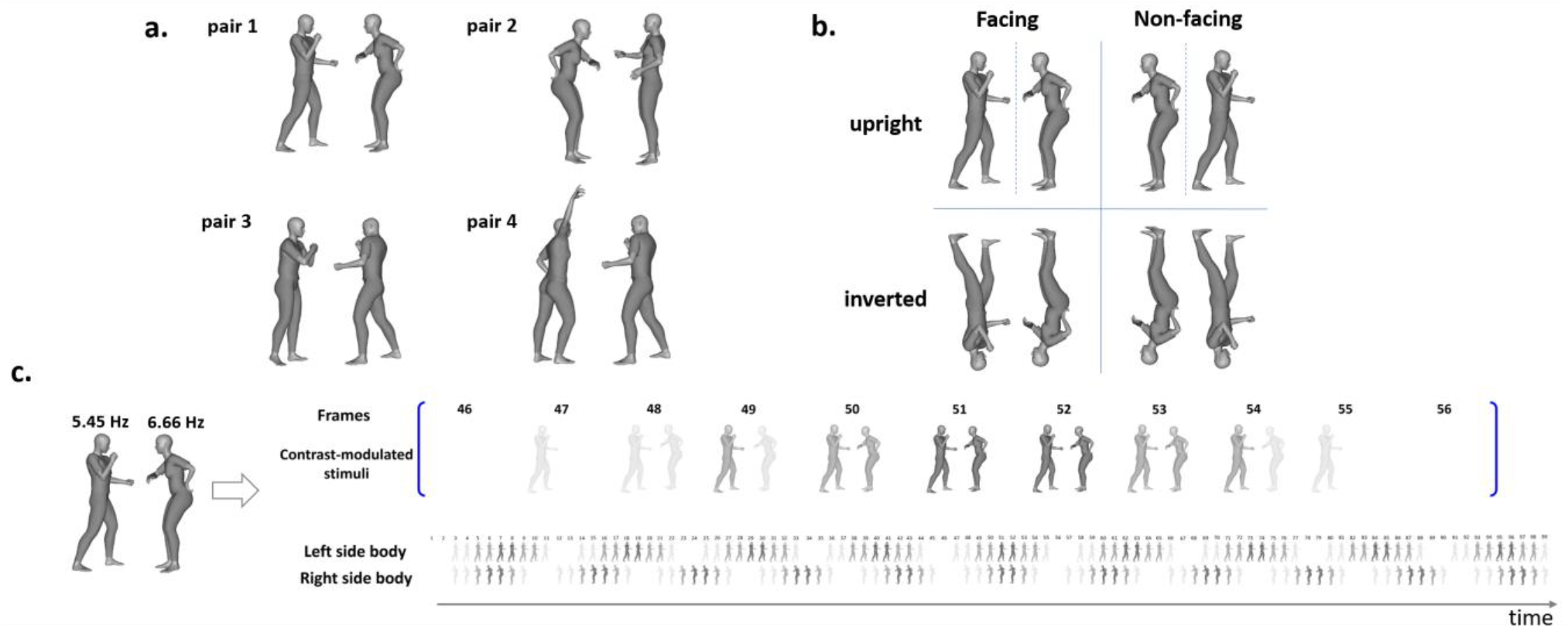
Illustration of stimuli, design and procedures for stimulus delivery. a) The four body dyads used in the study. b) Four variations of a dyad presented in a block. c) Illustration of the flickering presentation of the stimuli: the two bodies flickered at the frequencies F1 and F2, respectively. The contrast-modulated appearance of body images is shown over a cycle of 99 frames, corresponding to frequency Fd, at which both bodies appeared in full contrast. The blue brackets represent a zoomed-in illustration of the frames where the simultaneous appearance of the two bodies at their full contrast occurs (frames 51 and 52, as marked with the blue arrow).

For every (facing and non-facing) dyad, we created an inverted version by flipping the image by 180°. This yielded four variations of the same stimulus (i.e., upright facing, upright non-facing, inverted facing, and inverted non-facing), corresponding to the four experimental conditions of the study (Figure 1b).

### EEG Recordings

Brain activity was recorded using a 128-electrode EEG net (EGI, Eugene, USA), with a reference on the vertex. Recordings were continuously digitized at a sampling rate of 1000 Hz (net amp 400 system EGI, Eugene, USA). Participants were seated on a chair, one-meter away from a computer screen where the stimuli were presented. Single bodies on the screen had a mean visual angle of approximately 4.6° in height and 2.1° in width, and their inner edge were placed at 0.3° from the center. Dyads subtended a mean visual angle of 4.6° in height and 4.2° in width.

### Paradigm

The experiment consisted of four blocks. Each block included four trials of 70.91 seconds (~1.18 min) each, in which the same dyad was presented in one of the four experimental conditions (upright facing, upright non-facing, inverted facing, inverted non-facing). Trials were separated by a blank screen. After 20 seconds of blank screen, participants were free to start a new trial when they felt ready. Each block lasted about 6-8 min depending on the duration of the blank inter-trial intervals. The order of blocks and trials was randomized across participants. The experiment included a total of 16 trials (4 blocks, each including 4 trials). The total duration did not exceed 30 min.

Each trial consisted of fast periodic visual stimulation, where the two bodies flickered at frequency F1 of 6.6667 Hz (~ 6.66 Hz) and frequency F2 of 5.4545 Hz (~ 5.45 Hz), respectively (Figure 1c). Those frequencies were chosen so to prevent the overlap with the alpha band frequency (8–12 Hz) and, thus, maximize the signal-to-noise ratio of the evoked responses (Boremanse et al., 2013; Regan, 1989). That choice also took into account the refresh rate of the display (60Hz), to allow an integer number of frames for each cycle of stimulus presentation (60 Hz/6.6667 Hz ≈ 9 frames, and 60 Hz /5.4545 Hz ≈ 11 frames), preventing dropped frames or inaccuracies in the frequency of stimulus presentation. The contrast in the stimuli was sinusoidal-modulated (figure 1c). Stimuli were presented in cycles of 99 frames (9 × 11 = 99); in most frames, the two bodies were visible at a different contrast; they appeared together in full maximal contrast once during a cycle (at frames 51-52, see Figure 1c), corresponding to frequency Fd (dyad-related frequency) of 0.6061 Hz (60 Hz/99 ≈ 0.6061 Hz). Two input frequencies, F1 and F2, can give rise to non-linear interactions at frequencies nF1±mF2 (with n and m being integers; Zemon & Ratliff, 1984). Therefore, besides the response at Fd, the response at frequencies nF1±mF2 can reflect the interaction between processing stimulus 1 at F1, and stimulus 2 at F2. In our study, non-linear interaction frequencies corresponded to F1+F2 = 12.12 Hz and to F1-F2 = 1.21 Hz. We note that the latter response overlapped with the second harmonic of the response at Fd, F1-F2 = (n2-n1)Fd, where n1 (= 9) and n2 (= 11) are the number of frames corresponding to F1 and F2, respectively.

The oscillation frequency of a body (F1 or F2) and side of the screen (left or right) was kept constant for facing and non-facing dyads within a block, and was counterbalanced across blocks. Throughout a trial, a black cross was present at the center of the screen roughly aligned with the shoulders of the bodies (whether upright or inverted). The cross color changed from black to red for 200 msec, eight times at random intervals during the trial duration (i.e., 1.18 min). Participants were instructed to press the spacebar on a keyboard with their right hand when they detected the color change. This task was meant to maintain the participants’ vigilance and fixation at the center of the screen. Stimulus presentation, communication of triggers with the recording EEG system, and response collection were controlled with Psychtoolbox (Brainard, 1997) through MATLAB.

### Preprocessing

EEG recordings were band-pass filtered between 0.1 and 100 Hz using a zero-phase lag filter, and were further processed using the EEGLAB (Delorme & Makeig, 2004) and Brainstorm (Tadel et al., 2011) toolboxes in MATLAB. Recordings were segmented for the duration of the trials, based on the trigger marking the trial onset. In each trial, the first 3.3 seconds of the data (i.e., two cycles of stimulus presentation) were removed, taking into account the time needed for the entrainment to become effective on the brain activity. The analysis considered the following 66 seconds of the data for each trial, corresponding to 360 cycles for the stimulation frequency of 6.66 Hz, and 440 cycles for the stimulation frequency of 5.45 Hz, and 40 cycles of the dyad-presentation frequency 0.61 Hz. The final 1.6 seconds (i.e., one cycle of stimulus presentation) were not analyzed, to take into account the potential imprecision of boundary markers.

### Frequency domain analysis

In the periodic activity evoked by the periodic presentation of the stimuli, responses to the two individual bodies were frequency-tagged at F1 and F2, while responses to dyads were frequency-tagged at Fd. Time-series from each trial were transformed into the frequency domain using discrete Fourier transform, with a high frequency resolution 0.0152 Hz (1/66 sec = 0.0152 Hz), which allows studying the response at the frequencies of interest with high precision. For each resulting frequency spectrum, the Signal-to-Noise Ratio (SNR) was then computed as the amplitude at each frequency divided by the mean amplitude of the 80 neighboring frequencies (40 frequency bins on each side), after excluding the 4 immediately adjacent bins (Boremanse et al., 2013; Leleu et al., 2020). The SNR was then averaged across trials of the same experimental condition, separately for each participant. Mean SNR values were analyzed to determine the response to individual bodies and dyads, at the corresponding tagged frequencies.

### Statistical analyses and results

#### Grand-average identification of frequency-tagged responses

We first evaluated how many participants showed above-noise-level responses at the predefined frequencies, irrespective of experimental conditions and spatial distribution of the responses, to ascertain that our stimulation was effective in evoking frequency-tagged responses. To this aim, we averaged the SNR spectrum over all EEG sensors and trials for each participant (figure 2.a). Then, we compared the SNR at the frequencies F1 and F2 (6.66 Hz and 5.45 Hz), Fd (0.61 Hz) and their harmonics, and the second-order intermodulation responses at nF1±mF2, against the noise level (80 neighboring frequency bins excluding the immediately adjacent 4 bins), with a one-tailed *t*-test. This analysis showed significant responses at the main frequencies of interest F1, F2 and Fd, as well as at multiple harmonics of those frequencies (2 × F1/F2/Fd, 3 × F1/F2/Fd, 4 × F1/F2/Fd). Note that the even harmonics of the response at Fd also corresponded to harmonics of the intermodulation response F1-F2 (e.g., 2Fd = F1-F2). Other second-order intermodulation responses (i.e., F1+F2) were not significantly above noise-level in this whole-sensor analysis.

**Figure 2.**
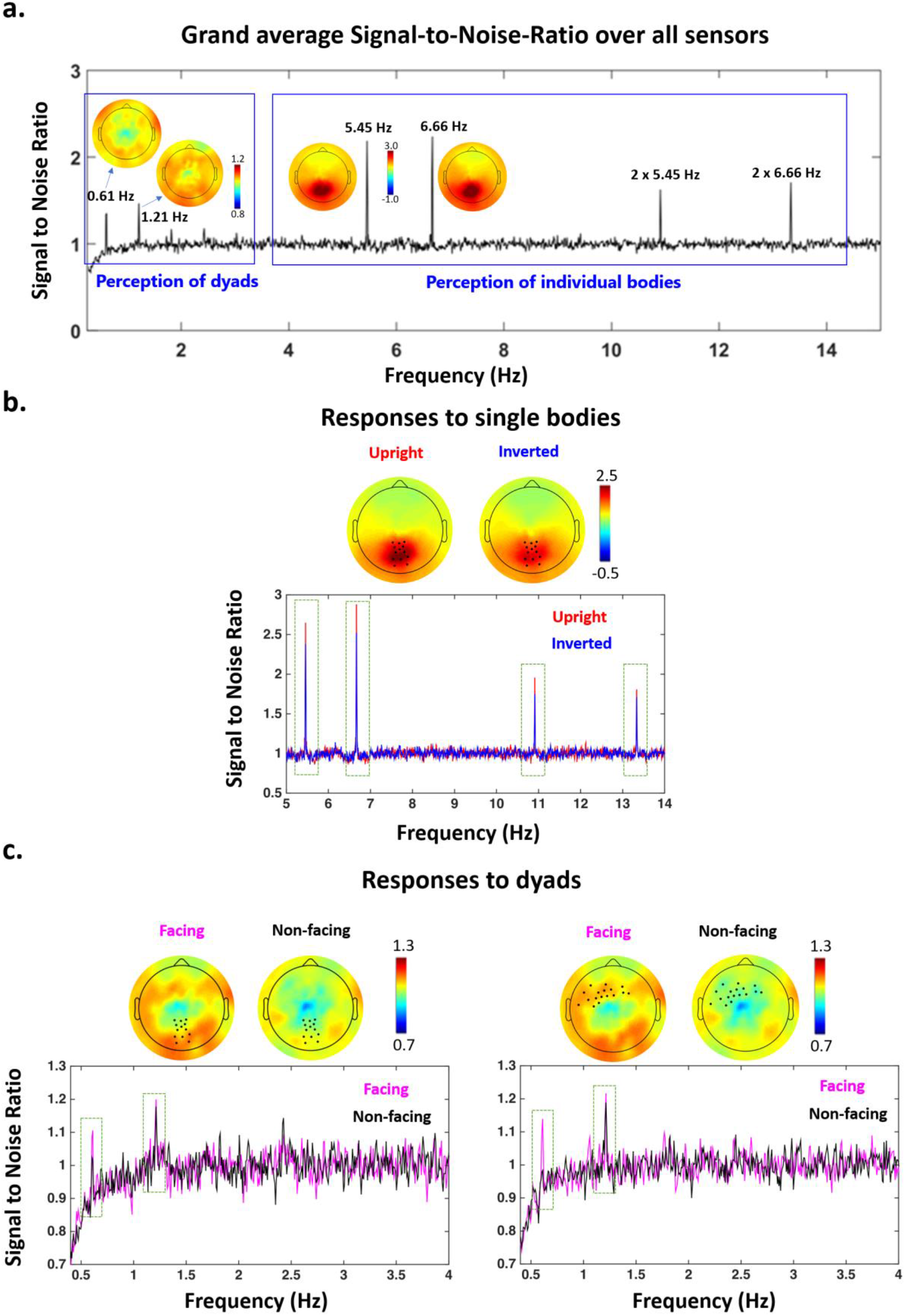
Frequency-tagged responses and their modulation depending on the experimental conditions. a) Grand average Signal-to-Noise-Ratio (SNR) over the group of participants, averaged over all EEG sensors. Peaks are visible at the stimulation frequencies corresponding to single bodies and dyads. Adjacent to the peak, the spatial distribution of responses is shown on a topographical map. b) Response to upright and inverted single bodies in a posterior cluster (body-related cluster) as highlighted on the topographical maps. Average SNR value within this cluster at frequencies F1 and F2 are highlighted in red for upright bodies and in blue for inverted bodies. c) Response to facing and non-facing dyads in the posterior body-related cluster of sensors (left), and in the anterior cluster pointed out by the analysis across all EEG sensors (right). These clusters are highlighted on the topographical maps and the average SNR values within each of them is plotted below the corresponding topographies, showing higher activations for facing dyads (in magenta) than for non-facing dyads (in black).

For both single-body and dyad-related responses, we constrained the analyses to the first two harmonics, since these responses were the most robust in 18 out of 20 participants and allowed keeping the same number of harmonics for both responses (statistical analysis of all harmonic responses is reported in Table 1). Thus, to study the response to single bodies, we averaged the SNR at 6.66 Hz and 2 × 6.66 Hz (body 1) and at 5.45 Hz and 2 × 5.45 Hz (body 2); to study the response to dyads, we averaged the SNR at 0.61 Hz and 2 × 0.61 Hz (1.21 Hz).

**Table 1.** Significance of the responses at stimulation frequencies of interest and their multiple harmonics as assessed on the grand-average SNR of all EEG sensors. Significant responses at the frequencies of interest correspond to SNR values higher that noise-level SNR. For responses to individual bodies, significance was tested for frequencies <50 Hz; responses to dyads, significance was tested for frequencies <5 Hz. P-values are Bonferroni corrected for the number of harmonics tested a frequency.

| Frequency | F1<br>6.66 Hz | 2 x F1<br>13.33 Hz | 3 x F1<br>20.00 Hz | 4 x F1<br>26.66 Hz | 5 x F1<br>33.33 Hz | 6 x F1<br>40.00 Hz | 7 x F1<br>46.66 Hz | 8 x F1<br>above 50 Hz | 9 x F1<br>above 50 Hz |
| --- | --- | --- | --- | --- | --- | --- | --- | --- | --- |
| Subjects (out of 20) with significant responses | 20 | 20 | 19 | 20 | 13 | 18 | 14 | -- | -- |
| Statistics | t = 8.7, p < 0.001 | t = 8.1, p < 0.001 | t = 5.3, p < 0.001 | t = 7.3, p < 0.001 | t = 2.3, p = 0.110 | t = 5.6, p < 0.001 | t = 2.5, p = 0.071 | -- | -- |
| Frequency | F2<br>5.45 Hz | 2 x F2<br>10.90 Hz | 3 x F2<br>16.36 Hz | 4 x F2<br>21.81 Hz | 5 x F2<br>27.27 Hz | 6 x F2<br>32.72 Hz | 7 x F2<br>38.18 Hz | 8 x F2<br>43.63 Hz | 9 x F2<br>49.09 Hz |
| Subjects (out of 20) with significant responses | 18 | 19 | 20 | 18 | 19 | 17 | 15 | 15 | 13 |
| Statistics | t = 6.2, p < 0.001 | t = 6.9, p < 0.001 | t = 6.3, p < 0.001 | t = 4.0, p = 0.003 | t = 4.5, p = 0.001 | t = 4.5, p = 0.001 | t = 3.7, p = 0.007 | t = 2.9, p = 0.04 | t = 2.6, p = 0.07 |
| Frequency | Fd<br>0.606 Hz | 2 x Fd<br>1.21 Hz | 3 x Fd<br>1.81 Hz | 4 x Fd<br>2.42 Hz | 5 x Fd<br>3.03 Hz | 6 x Fd<br>3.63 Hz | 7 x Fd<br>4.24 Hz | 8 x Fd<br>4.84 Hz | 9 x Fd<br>above 5Hz |
| Subjects (out of 20) with significant responses | 20 | 19 | 14 | 15 | 16 | 14 | 13 | 13 | -- |
| Statistics | t = 7.5, p < 0.001 | t = 6.4, p < 0.001 | t = 3.6, p = 0.007 | t = 3.2, p = 0.018 | t = 2.5, p = 0.083 | t = 2.0, p = 0.229 | t = 2.2, p = 0.163 | t = 1.2, p = 0.934 | -- |

Responses at F1, F2 and their harmonics were predominantly distributed over posterior electrodes with medial distribution. Responses at Fd and 2×Fd (=F1-F2) were more broadly distributed over the scalp, encompassing posterior and anterior areas (Figure 2).

The intermodulation response at F1+F2 was not significant in the first (whole-sensor) analysis. Since this response has often been described in similar frequency-tagging studies, we ran a secondary, less conservative, analysis, restricting our search space to the 20 posterior sensors extending from O1/O2 to T5/T6. We tested the SNR value at F1+F2 against the noise-level for each sensor using a one-tailed *t*-test. This analysis revealed a response at F1+F2 significantly above the noise level, on 3 adjacent electrodes (*t*(19) = 3.17; p=0.050, correcting for 20 multiple comparisons).

#### Definition of body-related response

Previous fMRI studies have reported that the encoding of spatial relations between multiple bodies begins in visual areas for body perception (Abassi & Papeo, 2020; Walbrin & Koldewyn, 2019). To follow up on those studies, we singled out visual activity for viewing bodies, using the contrast upright vs. inverted bodies at F1, F2, 2xF1 and 2xF2. Indeed, inverted bodies typically reduce or change the response to bodies in occipital visual areas (Brandman & Yovel, 2010; Brandman & Yovel, 2016; Minnebusch et al., 2009; Stekelenburg & de Gelder, 2004).

The comparison between SNR values for upright *vs.* inverted bodies was carried out for each sensor separately with a one-tail *t*-test. Neighboring sensors showing an effect with a *t*-value larger than a threshold corresponding to *p* < 0.05 (*t*(19) > 1.73) were clustered and tested for significance using nonparametric cluster-mass permutation test (Maris & Oostenveld, 2007), with 5000 random permutations of the condition labels on the original data. For each shuffled dataset, a *t*-test was carried out for each sensor separately. Neighboring sensors yielding above-threshold *t*-values were clustered together. The significance probability of the original clusters was computed as the number of times the shuffled data produced clusters with higher summed t values than the real data. This analysis revealed a posterior cluster of electrodes with significantly higher response to upright than inverted bodies (*p* = 0.03; Figure 2.b).

#### Effect of positioning and orientation of body dyads

In the first analysis, we targeted neural correlates of body perception and tested the hypothesis that body positioning affects the visual processing of bodies, as suggested by recent fMRI studies (Abassi & Papeo, 2020; Bellot et al., 2020; Walbrin & Koldewyn, 2019). In the cluster of electrodes that showed higher activity for upright *vs.* inverted single bodies (see above Section “*Definition of body-related response”*), we studied the effect of positioning (facing or non-facing) and/or orientation (upright or inverted) on the response to dyads. To this end, the mean SNR values for Fd and 2Fd (= F1-F2) over the predefined cluster were entered into a repeated-measures analysis of variance (ANOVA), with factors Positioning and Orientation. This analysis revealed a main effect of Positioning, *F*(1,19) = 5.5, *p* = 0.030, with higher activity for facing than for non-facing dyads (Figure 2.c), but no effect of Orientation, *F*(1,19) = 2.2, *p* = 0.152, or interaction, *F*(1,19) < 1, *n.s.*

In a second analysis without a priori hypothesis about the spatial distribution of the responses, we tested the effects of bodies positioning, orientation and their interaction on the SNR values, at each sensor, using repeated-measure ANOVAs. Neighboring sensors showing a main effect or an interaction with an *F* value higher than the threshold corresponding to *p* < 0.05 (*F*(1, 19) > 4.3) were considered as clusters. Clusters were tested for significance using nonparametric cluster-mass tests, with 5000 random permutations of the condition labels on the original data. For each shuffled dataset, an ANOVA was carried out, separately for each sensor. Neighboring sensors yielding above-threshold *F* values were clustered together. The significance of the original cluster was computed as the number of times the shuffled dataset produced clusters with higher summed *F* values than the real data. This analysis highlighted an anterior cluster, showing higher activity for facing than for non-facing dyads (cluster-mass permutation test: *p* = 0.03; Figure 2.c). We found no significant effect of orientation and no interaction between the two factors (all *p*s > 0.05).

Finally, we tested whether the average response at F1+F2 identified in the three posterior sensors was affected by positioning or orientation of bodies in the dyads. A repeated-measures ANOVA showed no effect of positioning (*F* < 0.1, *p* = 0.980), orientation (*F* = 4.0, *p* = 0.060), or interaction (*F* < 0.1, *p* = 0.875).

In summary, our analyses highlighted activations distributed over posterior and anterior sensors, suggesting an integrated representation of the two bodies in a face-to-face (relative to back-to-back) configuration.

#### Effect of single body direction

In this analysis, we assessed whether the difference between facing and non-facing dyads observed in posterior and anterior clusters, was due to the processing of the dyad as a whole (i.e., to the processing of the relation between two bodies), or could be rather accounted for by a difference in the visual treatment of each single body in the facing *vs.* non-facing condition. We reasoned that, in the latter case, the identified clusters should show different responses not only to dyads (i.e., at dyad-related frequencies Fd and 2Fd), but also to single bodies in the facing *vs.* non-facing condition (i.e., at the single-body related frequencies F1 and F2).

To this end, we tested the effect of positioning and orientation, separately for each cluster, considering the mean SNR values measured at F1 and F2. In the posterior cluster, this analysis confirmed the effect of orientation, *F*(1,19) = 18.5, *p* < 0.001 (see above Section *“Definition of body-related response”*), with no effect of positioning, *F*(1,19) = 0.2, *p* = 0.643, or interaction, *F*(1,19) = 0.2, *p* = 0.675. No effect reached significance in the anterior cluster (orientation: *F*(1,19) = 0.5, *p* = 0.498; positioning, *F*(1,19) = 0.7, *p* =0.417; interaction: F(1,19) = 3.2, *p* = 0.089). This analysis showed that the above effect of positioning of bodies in dyads could not be explained by the visuo-spatial features of single bodies such as the direction relative to central fixation (inward *vs.* outward).

## Discussion

Using a frequency-tagging EEG paradigm, we recorded frequency-separable responses to body dyads (at Fd and F1-F2) and to each single body that formed the dyad (at F1 and F2). We first identified neural response associated with perception of single bodies at F1 and F2, using the contrast of upright *vs.* inverted bodies (Stekelenburg & de Gelder, 2004). This analysis revealed a posterior cluster, compatible with the fMRI activity evoked in the occipitotemporal cortex, by viewing canonical upright (vs. inverted) bodies (Brandman & Yovel, 2010; Brandman & Yovel, 2016). Within that cluster, we tested the hypothesis that the neural correlates involved in body perception are sensitive to the relative positioning of multiple bodies in a visual scene. In line with previous studies (Abassi & Papeo, 2020; Bellot et al., 2020; Walbrin & Koldewyn, 2019), we found an effect of positioning, with larger response to facing than to non-facing dyads at Fd and F1-F2. Extended to the whole scalp (i.e., all the sensors), our analysis revealed another cluster with anterior distribution, showing larger response to facing than to non-facing dyads.

Unlike the responses evoked at the dyad-related frequencies, the responses evoked by single bodies did not vary depending on whether a body was part of a facing or a non-facing dyad. Thus, the effect of positioning at Fd and F1-F2 captured the response to the whole dyadic stimulus, rather than a change in the response to a single body seen in a facing vs. a non-facing dyadic context. In other words, the effect reported here appears to reflect encoding of relative positioning (i.e., the positioning of a body relative to another), rather than the absolute body positioning (i.e., body orientation directed toward or away from the center).

While the neural response to single bodies, at F1 and F2, was reduced when bodies were presented upside-down (see also Brandman & Yovel 2010; Brandman & Yovel, 2016; Minnebusch, Suchan & Daum, 2008; Stekelenburg & De Gelder, 2004), we did not see any effect of inversion on the neural response to dyads (i.e., at Fd 0.61 Hz and F1-F2). Although multiple bodies have also been shown to be susceptible to the effect of inversion (Abassi & Papeo, 2019; Papeo et al., 2017), it is possible that such effect is visible at high stimulation frequencies (i.e., the frequency of single-body presentation) but not at slower stimulation frequencies (i.e., the frequency of dyad presentation). Moreover, a difference between upright and inverted dyads might not occur in the frequency domain, but in the time domain (responses might have different shape in the time course). Future studies should address this question using, for example, event-related designs.

Irrespective of the orientation, the response to dyads was effectively tagged at two frequencies, possibly reflecting different processes: Fd (0.61 Hz), corresponding to the periodic simultaneous appearance of two bodies in full contrast, and 1.21 Hz, corresponding to the second harmonic of Fd (2 × 0.61 Hz = 1.21 Hz), and the nonlinear interaction between the two input frequencies (F1-F2 = 1.21 Hz; Zemon & Ratliff, 1984). As one can appreciate from figure 2.a, the response at 0.61 Hz was comparable in strength with the response at 1.21 Hz. Since harmonics typically have lower amplitudes than the main response, it is less likely that the response at 1.21 Hz only reflected the harmonic of the response at Fd. This raises the possibility that the response at 1.21 Hz resulted from both the second harmonic of the response to the periodic visual stimulation (i.e., the periodicity of dyad presentation at Fd), and the nonlinear interaction of the response to the two stimulation frequencies (F1-F2) (Alp et al., 2016; Boremanse et al., 2013). Future studies should be designed to distinguish between the visual responses to the whole scene, (i.e., at Fd), and the nonlinear interactions between the parts that give rise to the whole scene, as marked in the emergent intermodulation responses at nF1±mF2.

The response to dyads at other intermodulation frequencies such as F1+F2, was clearly weaker than the response at Fd/F1-F2 in our study, as it was only observed when restricting the analysis to posterior sensors, in three adjacent sensors. The response at F1+F2, as weak as it might be, further supports the involvement of integrative processes that arise from nonlinear interaction of the main (single body) responses.

Frequency-tagged responses at the dyad-related frequencies Fd and F1-F2 marked the neural representation of the two bodies together. Note that, the two bodies were present on the screen in most frames of a trial, although at a varying contrast level. That is, participants experienced the presence of two flickering bodies throughout the trial and were not aware that the bodies were only occasionally (i.e., at Fd) shown together in full view. This feature of the paradigm implies that the response to dyads at Fd captured a process that was spontaneous –if not automatic– given certain input characteristics, and independent of an explicit goal or task (Fodor, 1983); most likely a perceptual process. On this background, the larger response to dyads of facing (vs. non-facing) bodies suggests the recruitment of additional processing, distributed over posterior and anterior areas, beyond the processing of multiple single bodies that were identical in facing and non-facing dyads. Although previous studies have primarily focused on the effect of spatial relations between bodies in posterior (visual) areas, increased frontal activity for facing dyads has also been observed (Abassi & Papeo, 2020; Bellot et al., 2020). We note that, here, posterior activity was higher for facing dyads, but was also present for non-facing dyads; instead anterior activity was observed for facing dyads, but was virtually absent for non-facing dyads at Fd. Higher posterior (i.e., visual cortex) activity for facing dyads has been interpreted as an effect of the visual enhancement of multiple-body configurations that cue interaction (Papeo, 2020). Less clear is the role of frontal activity in this processing. The selectivity of anterior activity observed here for facing dyads, opens to the hypothesis that it reflects a binding process triggered when two bodies are combined into a unitary representation.

We interpret the facing *vs.* non-facing dyad effect at Fd and F1-F2 in the spirit of previous studies, where frequency-tagging paradigms were used to mark the integrative processing that mediates grouping of multiple parts into a whole, such as geometrical elements into a new object representation (e.g., the Kanizsa triangle; Aissani et al., 2011; Alp et al., 2016; Gundlach & Müller, 2013), or face-halves in a face (Boremanse et al., 2013). Those studies demonstrated that spatial relations between parts are key to trigger the integration captured by the enhanced response to a canonical *vs.* noncanonical configuration. Thus, for example, two face halves evoked increased response at the integration frequency when aligned but not when misaligned (Boremanse et al., 2013). By analogy, in the current study, two bodies increased the response at the alleged integration frequency, when they appeared simultaneously and facing toward –but not away from– each other. Adding to the extant literature, our results show that grouping can account not only for object formation through binding of disconnected, but spatially organized, parts (Aissani et al., 2011; Alp et al., 2016; Boremanse et al., 2013; Gundlach & Müller, 2013), but also for the formation of scenes (or events) by binding multiple disconnected objects (here, bodies) together. The binding mechanism that would underlie the increased response to facing dyads may account for behavioral and neural effects showing attentional/perceptual advantages for sets of objects in spatial configuration that cues interaction (e.g., face-to-face bodies; see Papeo, 2020), a common function or usage (e.g., a pen over a notebook; see Kubilius et al., 2015), or a coherent scene (e.g., a lamp above the table; see Kaiser et al., 2019). It remains unknown whether a single domain-general mechanism accounts for binding of objects of different categories and according to different types of relations (e.g., bodily/social interaction, statistical regularity, semantic relatedness). In other words, whether the effect described here for facing body dyads is specific to bodies, or it applies to other object sets that can form coherent scenes (e.g., a lamp above a table) or functional groups (e.g., a pen above a notebook), remains an outstanding question for future research.

In conclusion, by contrasting the response to facing *vs.* non-facing dyads involving the same individual bodies, we were able to disentangle objective neural responses to parts (individual body) and to the whole scene (the dyad). The response to facing (*vs.* non-facing) dyads described here echoes neural effects that have been consistently associated with perceptual grouping (also called visual binding), and suggests that the face-to-face body positioning can trigger a new representation that is more than the sum of the constituent parts. The representational content of the composite representation emerging from two facing bodies remains a fascinating question for further research. By extending the frequency-tagging paradigm to a new class of stimuli (i.e., multiple-body stimuli), our results contribute to characterizing grouping as a general mechanism that could account not only for the representation of objects from parts, but also for the representation of scenes from objects or events from social entities.

## Acknowledgments

This work was supported by ANR grant awarded to J-R. H. (Project: TACTIC, Grant Agreement ANR-16-CE28-0006) and European Research Council Starting Grant awarded to L.P. (Project: THEMPO, Grant Agreement 758473).

